# Infestation and growth of *Tribolium confusum* (Coleoptera: Tenebrionidae) on two varieties of local durum wheat: nutritional value and varietal susceptibility

**DOI:** 10.1101/2022.05.11.491516

**Authors:** Haffari Faouzia, Boualem Malika, Bergheul Saida, Merzoug Aicha

## Abstract

*Tribolium confusum* is a pest of stored produce on many cereals. This study was planned to see quantitative losses and feelings of the equipped grains owed to the infestation by the red genus Tribolium on 2 forms of native durum in Algeria. Selection Puts tire chains S am cultivated within the region of Adrar, underneath associate arid climate; and also the existent selection Vitron of a production of the region of Ain Témounchent underneath a dry Mediterranean climate. Study was accomplished during two months to determine the weight loss of grains, damage caused by *T. confusum* and the influence of two varieties of hard wheat on growth and reproduction of this vandal. Damage of the couples (5, 10, 15, 20) on grains showed significant difference. A similar tendency was noted for the weight loss of grains Put snow chains S; while Vitron noticed not significant losses. Also, an increase in the consumption of proteins, lipids and starch of grains was noticed with the increase of the length of stocking. The appearance of generations was variable according to the level of density and the variety; nevertheless, it was generally highest on Vitron and low on Chain S. The results show that the quantitative and qualitative losses of stored grains are linked to the storage time, the chemical composition of the grains and the susceptibility to insects of the stored products. These results can be integrated into food safety management to ensure the quality of stored wheat and their stored food derivatives.

## Introduction

Pests and diseases are the major obstacle of man to agricultural products, where pests are the principal competitors of man for the resources generated by agriculture (HAGSTRUM & SUBRAMANYAM 2009; OLIVEIRA *et al*. 2014). Preserving the productivity of crops, protective them from harm caused by weeds, animal pests and pathogens is additionally a necessary condition for providing decent food and put in decent amount and quality (OERKE & DEHNE 2004). The damage caused by these pests is a major factor in reducing the productivity of many crops, either in the field or later in storage (OERKE 2006).

Stored crops represents an important part of the economy and human nutrition. Cereals are the most frequently stored products thanks to their low water content (GARCÍA-LARA & SALDIVAR 2015). Cereal production is the basis of global food security (MESTERHÁZY *et al*. 2020). In Algeria, the 2020 cereal production is estimated at an above average level of 5.6 million tons, about 8% below the record level of 2019 due to the amount and distribution of rainfall, dry weather conditions (FAO 2021).

Durum wheat is the third cereal cultivated in the world, it is adapted to more diverse environments than bread wheat, and it performs very well in semi-arid regions (KADKOL & SISSONS 2015). The cultivation of durum wheat (*Triticum turgidum* L. var. Durum Desf.) is used for a wide range of food products around the world (ALZUWAID *et al*., 2020).

Stored wheat is subject to attacks from different insect species which reduce the quality of infested products, causing quantitative and qualitative problems (SAAD *et al*. 2018; TADDESE *et al*. 2020). 1,660 insects species including 600 beetles species were reported to be associated with stored products (HAGSTRUM & PHILLIPS 2017; HAGSTRUM & SUBRAMANYAM 2009). FAO (2019) estimates that each year between 20% and 40% of global agricultural production is lost to pests, the total potential global loss due to pests are around 50% for wheat (GARCÍA-LARA & SALDIVAR 2015). Post-harvest losses caused by pests represent 10-36% of grain weight worldwide (GITONGA *et al*. 2013; HAGSTRUM & PHILLIPS 2017).

Among these storage predators, *Tribolium confusum* (Duval) (Coleoptera: Tenebrionidae) is classified as a secondary colonizer because it can develop more easily on grains broken or already infested by primary colonizers. ATHANASSIOU & ARTHUR (2018) AND HERNANDEZ NOPSA *et al*. (2015) have highlighted the serious hazard to health that the carcinogenic effect of benzo-quinones, secreted by this pest of stored foodstuffs can induce.

Crop losses caused by pests were calculated for all crops on a regional basis; depending on the intensity of crop production and production conditions (OERKE & DEHNE 2004). The objective of this research is to know the behavior of *Tribolium confusum* according to two varieties of durum wheat. Therefore, this study may provide information on the biology and development of *Tribolium confusum* on the two local durum wheat varieties necessary in the investigation of control strategy.

## Materials and methods

### Plant material

Samples are collected from the Union of the agricultural Cooperative of Mostaganem and therefore the Cooperative of Grain and Pulses of Adrar (Algeria). 2 varieties were collected of the assembly of the year 2018/2019 to be famed, place chain on them S and Vitron R1 cultivated within the region of Adrar, below associate arid climate; and therefore the existent selection Vitron of a production of the region of Ain Témounchent below a dry Mediterranean climate

### Animal material

The people of *confusum* genus *Tribolium* employed in the course of this study were collected within the organism of stocking UCA Mostaganem. The agriculture of this type is created as: 250 g of grains of *Triticum durum* already troubled by genus *Tribolium* red, is adults, one in every of eggs or of grubs took a sample haphazardly and place in glassy cylindrical jars of nine.5 cm, diameter and eleven.5 cm high, coated by a textile screen with fine sew to permit the ventilation of insects and preventing them from going out. thirty not sexual grown-up insects of *T. confusum* were separated introduced within the glassy jars. Agriculture takes place darkly during a etude regulated during a temperature of twenty seven ± 3°C and a relating wetness of 70±5 you have to enable one develop favorable to the insect. Having assured pairing and egg laying of eggs, the adults’ area unit eliminated and once thirty two days, the primary emergence seem. The appeared adults are eliminated, then they add 10g of hard wheat to jars containing the new generations to assure their good development the previous same conditions. 28 - 37 days afterwards, the second emergencies appear, the same method is used for the third emergence. After 10 days, they recover the young emergent adults, who will be used for the next tests.

#### Method of Study

Two verity of *Triticum durum* chosen for the check, a 1000 grains taken at random; their weights are determined. Four countless *Triticum durum* were overrun with adults of *T. confusum* (5, 10, 15 and 20 pairs), in 5 repetitions; and incubated for 60 days. That we tend to select the most length of sixty days for our check, this by relating the work meted out by UAC-Mostaganem, that mentions that *Triticum durum* needs a most storage amount of two months. Every day, the load and coincident rising and dead people are determined at a mean temperature and a relative humidity of 27 ± 3 °C and 70 ± 5%.

#### Parameters studied

##### Analyses physicochimique

analyses physic chemical of *Triticum durum* were determined according to norms following: specific weight is determined according to norm NF V03-719 (1988); water content: NA: 1.1.32 (1990) and JO n ° 08 (2013); ash: ISO 2171 (2007) and JO n ° 35 (2006); protein content: NF V03-05 AFNOR, by the determination of the content of complete nitrogen according to the method of KJELDAHL (1973); lipid content: NF. ISO 734 – 1 (2000).

##### Grain loss rate (%)

According to SEMPLE *et al*. (1992), the proportion of grains attacked by *T. confusum* was calculated using the following equation:

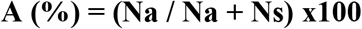

Number of grains attacked (Na); Number of healthy grains (Ns)

##### Weight loss (%)

As reportable by HUIGNARD (1985), insect harm in stocks isn’t perpetually simple to assess. Totally different ways have planned to work out the load losses. The foremost ordinarily used area unit the determination of attack and weight loss rates; to try to do this, the grains separated into 2 batches, one in all N healthy grains (Ns), and also the different of N attacked grains (Na). The grain weight loss estimates victimization the equation below:

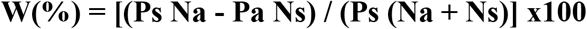

- Weight loss rate (W), Number of grains attacked (Na); Number of healthy grains (Ns); Weight of the attacked grains (Pa); Healthy grain weight (Ps).

##### Effect of varieties on development time of *T. confusum*

The dates of infestation and emergence were recorded so as to work out the mean length of development (Dmd) in step with following expression:

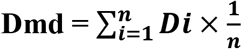

Dmd: average development time; Di: development time of an individual i on a given variety and n = total number of individuals.

##### Growth rate (Gr)

Development of *T. confusum* on each sort of tested native durum wheat is calculable per expression below:

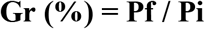

Pf = final population, Pi = initial population.

##### Dobie Index (ID)

He permits to characterize the extent of feelings of a range to a damaging of providers. For a given selection, a lot of indication is well remarked and therefore a lot of selection is sensitive (DOBIE 1977):

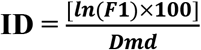

F1 = Number of individuals of the first generation; Dmd: Average development time.

##### Germination power

Of every many takes a look at, one took a sample of a hundred grains then at place within the cotton soaked in water in one limp with Kneading while not cowl. At the top of week, the germinated grains are going to be enumerated for each heap. The speed of germination is given by following expression:

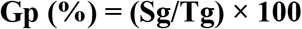

**Sg: number of sprouted grains Tg: total number of grains tested**

### Statistical analysis

Data on the percentage of damaged kernels, weight loss and growth rate of the two reconstituted native *Triticum durum* varieties were subjected to analysis of variance (ANOVA) and Tukey’s test at P <0.05. All statistical analyzes were performed with SPSS 26 software.

## Results

### Chain S and Vitron physicochemical analyzes

These analyses were applied to see the content of proteins, fats, water and ash in every kind of native *Triticum durum* before being employed during this study. This step was accomplished to estimate the technological quality of native *Triticum durum* (Table 1).

**TABLE 1.** Physicochemical parameters of wheat varieties studied. The values are means ± SD, Sw: specific weight, H: water content, Dm: dry matter, A: ash, Pr: protein content, Lp: lipid content.

|  | Parameters |  |  |  |  |  |
| --- | --- | --- | --- | --- | --- | --- |
| varieties | Sw (Kg/hl) | H (%) | Dm (%) | A (%) | Pr (%) | Lp (%) |
| Chain s | 78.6±0.06 | 15.28±0.27 | 84.72±3.63 | 1.18±0.20 | 15.96±0.17 | 1.44±0.08 |
| Vitron | 77±0.14 | 14.16±0.99 | 85.84±0.81 | 1.71±0.56 | 14.25±0.61 | 1.36±0.13 |

### Grain damage and weight loss caused by *T. confusum*

The percentage of damage caused by *T. confusum* within the grains of the provided wheat was significant for each variety: Chain S and Vitron (F = 20.418, P <0.000; F = 191.865, P <0.000, respectively). When comparing different tries, Vitron observed averages of 28.02 ± 1.08 for the twenty couples evaluated by the foremost well said. In effect, the amount of couples (F = 14,989; P <0,000) hampered the pace of weight loss of the Vitron grains provided (Table 2). Mile, multivariate study of weight loss of Chain S provided was not significant (F = 2.574; P <0.090) (Table 3).

**TABLE 2.** Comparison of damage caused by T. confusum within the grains of native durum wheat (Chain S and Vitron) consistent with the amount of introduction of the couples over 2 months of stocking. Letters (abc) estimated degrees of signification at P <0.05 (Tukey’s test) Values are means ± SD

| Varieties | 5CP | 10CP | 15CP | 20CP |
| --- | --- | --- | --- | --- |
| Chaine S | 6.44± 0.10<sup>a</sup> | 8.78 ±0.17<sup>a</sup> | 11.98± 0.09<sup>bc</sup> | 13.2 ±0.18<sup>c</sup> |
| Vitron | 12.72 ±0.15<sup>a</sup> | 19.64± 0.87<sup>b</sup> | 25.66 ±1.33<sup>b</sup> | 28.02 ±1.08<sup>bc</sup> |

**TABLE 3.** Comparison of the medium rate of weight loss of the grains of Chain S and Vitron caused by T. confusum in grains consistent with the degree of introduction of the couples over 2 months of stocking. Letters (ab) estimated degrees of signification at P < 0.05 (Tukey’s test), Values are means ± SD

| Varieties | 5CP | 10CP | 15CP | 20CP |
| --- | --- | --- | --- | --- |
| Chain S | 3.06 ±0.04<sup>a</sup> | 5.33±0.01<sup>a</sup> | 7.56 ±0.08<sup>a</sup> | 8.28 ±0.06<sup>a</sup> |
| Vitron | 5.46 ±0.15<sup>a</sup> | 7.24 ±0.61<sup>a</sup> | 9.15± 1.36<sup>ab</sup> | 13.81± 0.62<sup>b</sup> |

### Susceptibility of durum wheat varieties to infestation by *T. confusum*

The development cycle of *T. confusum* has an average duration of 29-36 days. The overall ANOVA was non-significant for the varieties Chain S and Vitron (F = 0.684, P <0.575; F = 0.484, P <0.698, respectively). *T. confusum* developed more rapidly on the variety Vitron in contrast to the variety Chain S (Table4). The rate of population increase of *T. confusum* was not significant for both varieties, but Vitron showed the highest rates of population increase of *T. confusum* recorded with an average of 2.58 ± 0.32 for the 15 pairs compared to Chain S. Both durum wheat varieties tested were susceptible to *T. confusum* attack, with a non-significant Dobie index. Chain S had the lowest Dobie index (F = 0.809, P <0.509). while Vitron has the highest Dobie index (F = 1.819, P <0.184) (Table 5).

**TABLE 4.**
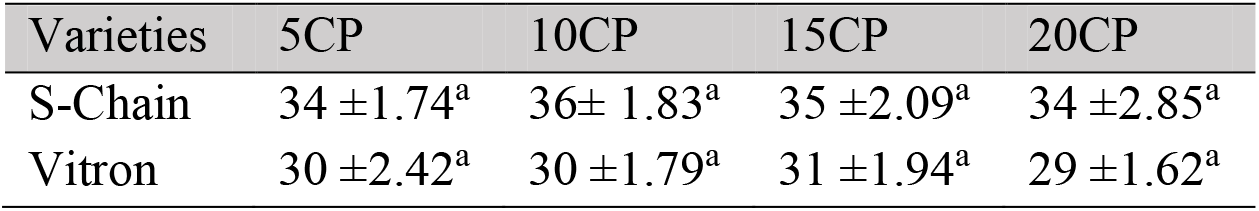
Length of development of *T. confusum* non heritable on each form of native *Triticum durum*. Letter (a) estimated degrees of signification at P < 0.05 (Tukey’s test), Values are means ± SD

| Varieties | 5CP | 10CP | 15CP | 20CP |
| --- | --- | --- | --- | --- |
| S-Chain | 34 ±1.74<sup>a</sup> | 36± 1.83<sup>a</sup> | 35 ±2.09<sup>a</sup> | 34 ±2.85<sup>a</sup> |
| Vitron | 30 ±2.42<sup>a</sup> | 30 ±1.79<sup>a</sup> | 31 ±1.94<sup>a</sup> | 29 ±1.62<sup>a</sup> |

**TABLE 5.**
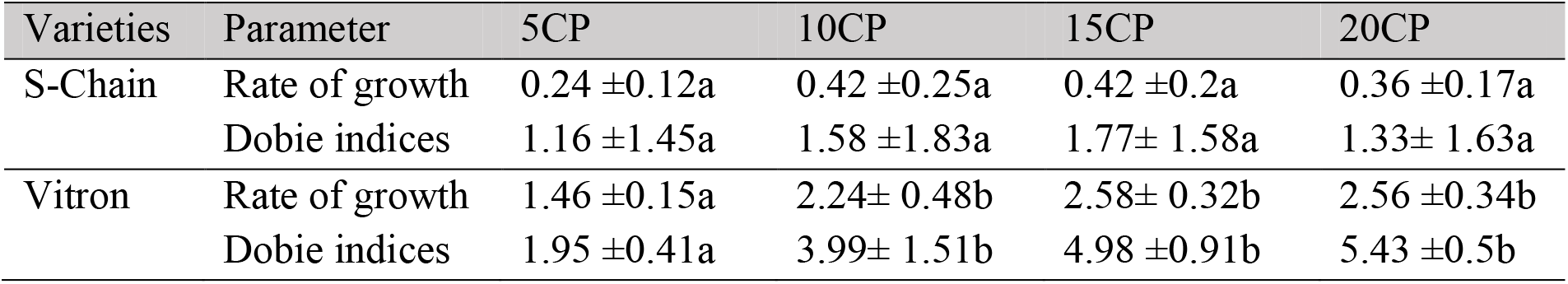
Rate of increase of *T. confusum* and Indications of Dobie recorded on each style of native *Triticum durum*. Letters (ab) estimated degrees of signification at P < 0.05 (Tukey’s test), Values are means ± SD

After the sexual activity of the adults of *T. confusum*, egg laying and egg abundance varied between the number of pairs studied and between the two wheat varieties tested. The ANOVA showed that the number of eggs recorded on Vitron was statistically significant (F = 14,280, P <0.000), while statistical analysis revealed non-significance for the same parameter studied on Chain S (F = 1,875, P <0.175). The abundance of larvae and adults counted showed a significant difference on Vitron (F = 15.518, P <0.000; F = 23.077, P <0.000, respectively). On the other hand, it is not significant on Chain S (F = 0.656, P <0.591; F = 643, P <0.598, larvae and adult respectively).

The mortality results of T. confusum during this study show significance on both varieties studied (Chain S, Vitron; F = 30.188, P <0.000; F = 9.812, P <0.001, respectively). Furthermore, the sex ratio analysis of T. confusum was not significant on Chain S and Vitron (F = 1.647, P <0.218; F = 1.957, P <0.161, respectively) (Table 6).

**TABLE 6.** Medium abundance of eggs, grubs, adults, mortality and sex-ratio of T. confusum below the influence of 2 styles of native arduous wheat; consistent with the degree of introduction of the couples over 2 months of stocking. Letters (abc) estimated degrees of signification at P < 0.05 (Tukey’s test), Values are means ± SD

| Varieties | Torques / Parameters | 5CP | 10CP | 15CP | 20CP |
| --- | --- | --- | --- | --- | --- |
| S-Chain | Egg | 2.4 ± 1.35a | 5.8 ± 3.48a | 7 ± 2.61a | 3.81 ± 3.44a |
| | Larva | 1.2 ± 0.98a | 3 ± 1.52a | 3.2 ± 2.04a | 2.4 ± 0.49a |
| | Adult | 0.6 ± 0.1a | 1.4 ± 1.49a | 1.8 ± 1.6a | 1.6 ± 0.2a |
| | Mortality | 7.4 ± 1.02a | 14 ± 1.79b | 21.8 ± 2.04c | 26.4 ± 5.39c |
| | Sex ratio | 0.2 ± 0a | 0.9 ± 0.11a | 0.8 ± 0.17a | 0.17 ± 0.11a |
| Vitron | Egg | 51 ± 27.13a | 88.6 ± 21.45ab | 118 ± 13.44bc | 143.4 ± 19.81c |
|  | Larva | 35.6 ± 21.63a | 62 ± 23.23ab | 95.4 ± 13.86bc | 117.6 ± 12.19c |
|  | Adult | 22.8 ± 10.59a | 36.2 ± 14.34ab | 69.4 ± 11.82bc | 96.4 ± 17.5c |
|  | Mortality | 1.8 ± 0.17a | 3.4 ± 1.02ab | 5.6 ± 2.15bc | 7.6 ± 1.85c |
|  | Sex ratio | 1.2 ± 0.5a | 0.83 ± 0.23a | 1.13 ± 0.5b | 1.35 ± 0.07b |

An analysis of correlation was additionally performed in SPSS. 26 by combination of all levels of relative density of the couples, production of derivation, feeding damages, and loss of paid were all arduous corrects absolutely some with the others (Table7).

**TABLE 7.** Correlation of the harm of grains, loss of paid and development of T. confusum combined for levels of the couple, r given stocks, P < 0,01 for all.

| varieties |  | Damage | weight loss | Growth |
| --- | --- | --- | --- | --- |
| S-Chain | Damage | | $r = 0.359, p = 0.120$ | $r = 0.084, p = 0.726$ |
| | weight loss | $r = 0.359, p = 0.120$ | | $r = 0.833, p < 0.000$ |
| | Growth | $r = 0.084, p = 0.726$ | $r = 0.833, p < 0.000$ | |
| Vitron | Damage | | $r = 0.223, p = 0.374$ | $r = 0.367, p = 0.135$ |
| | weight loss | $r = 0.223, p = 0.374$ | | $r = 0.763, p < 0.000$ |
| | Growth | $r = 0.367, p = 0.135$ | $r = 0.763, p < 0.000$ | |

### Germination power

Results of the effect of T. confusum couple abundance on the germ inability of durum wheat are shown in Figure 01. The readings recorded for this test highlighted the impact of the number of T. confusum pairs on the germination rate of each durum wheat variety tested. Indeed, the increase in pairs caused a decrease in germination rate. The ANOVA showed a significant difference in germination rate for the two varieties studied (Chain s, Vitron: F = 6.284, P <0.017; F = 64.672, P <0.000, respectively).

**Figure 1.**
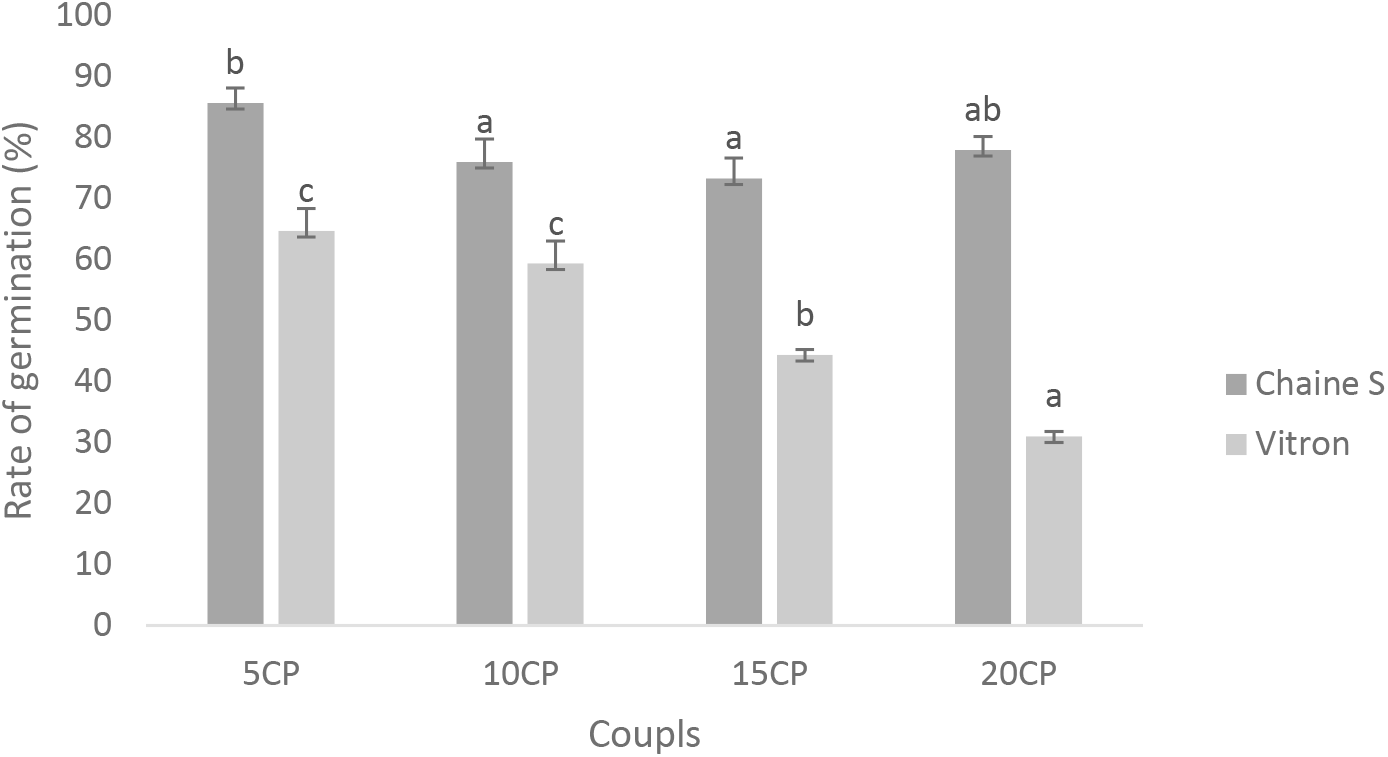
Rate of germination (Mean and Standard deviation)grains of 2 varieties tested in relevance the couples of *T. confusum*. Totally different letters imply vital distinction between varieties – couples(CP) (factorial multivariate analysis during a way; Tukey in = 0,05).

## Discussion

In Algeria, damage caused in grain will attain 64 %; these losses are owed to activities of insects, micro-organisms and animals. As well as poor handling and physical and chemical changes, all of which are interrelated. In the case of severe infestation by these pests, there is a significant loss of weight and a deterioration in quality (FAO 2019).

Results obtained above show that post-harvest insect pests cause heavy losses in stored grain. Also, a difference in susceptibility of the local durum wheat varieties studied to *T. confusum* is demonstrated. CHUCK-HERNÁNDEZ *et a*l. (2013); MAKATE (2010); SIWALE *et al*. (2010); TOEWS *et al*. (2003); WAONGO *et al*. (2018), report that the susceptibility of cereal varieties to storage pest attacks is related to the chemical and/or physical compositions of the grains.

Our study showed a non-significant effect of physico-chemical characteristics on Chain S and Vitron such as water, protein and starch content. The internal chemical composition may also play an important role in the tolerance of insects to stored products (ARTHUR *et al*. 2020). ARTHUR *et al*. (2012) and PERIŠIĆ *et al*. (2021) note that insect damage is relatively related to grain sensitivity and nutritional quality.

During this two-month study, we observed a very high rate of weight loss on the highest density of pairs, on both varieties tested. GONZÁLEZ-TORRALBA *et a*. (2013) have shown that weight losses of wheat grains can reach rates of 6-8% during a five month storage period. Increased storage time with favorable temperature and humidity can ensure mould growth as well as insect infestation resulting in significant grain weight loss (SUJITHA *et al*. 2018; TADDESE *et al*. 2020; TEFERA 2012). Preliminary studies have articulated that insect losses may be influenced by wheat varieties since insect multiplication could also be a lot of lower on some wheat varieties than on others (THRONE *et al*. 2002).

Life cycle from egg to adult in laboratory-reared strains of *T. confusum* can be short, 4 weeks under optimal conditions of food, temperature and humidity. Adults can live for more than 3 years, although a life span of 1-6 months is more typical in the laboratory (PAI 2009; SCHEFF & ARTHUR 2018). The growth cycle is 20 days at 38°C with an optimum of 30 days at 32°C, the population can be multiplied by 60 on optimal food in 28 days at 70% relative humidity (BELL 2011). Varieties with a higher susceptibility index produce a lot of offspring and vice versa, varieties with a lower susceptibility index produce a minimal number of offspring (ASTUTI *et al*. 2013; FAJARWATI *et al*. 2019; SIWALE *et al*. 2010). Corresponds to our study, we have a tendency to distinguish a development cycle of *T. confusum* of 29 to 36 days at temperature of 27 ± 3 ° C and a relative humidity of 70 ± 5%.

the insect can cause a substantial loss of germination on seeds since the larvae feed particularly on the germ (STEJSKAL *et al*. 2015). Any kind of damage to the grain can cause a significant reduction in its ability to produce a healthy plant (PÉREZ *et al*. 2008). Damage accelerates the loss of nutrients during the initial stages of seed germination and the seeds fail to develop into normal plants (KALSA *et al*. 2019). Our result shows that the germination of grains reduces significantly with the increase of the specific gravity of the couples of *T. confusum*.

Our results show that durum wheat varieties varied in their tolerance to T. confusum. Chain S is a T. confusum resistant variety compared to Vitron. The latter has a higher ash content which makes it more susceptible to *T. confusum* infestation. The two-month reproduction trial on the two varieties tested showed a variation in the number of pairs studied, with an increase in the population in the next generation. In addition, grain weight loss and damage on both varieties were related to storage time and chemical composition of the nutrient substrate. The types of wheat tested were an influential factor in the development of the T. confusum pest. It can be concluded from this study that genetics, environmental factors, such as temperature, humidity and food also influence the development of this pest in stored commodities.

## Supporting information

figure jpg

## Acknowledgment

We would like to thank Dr. MAAMAR M. of the Union of Agricultural Cooperative of Mostaganem (UCA) and Dr. HAMADAT K. of the Cooperative of Cereals and Pulses of Adrar (CCP) who provided us with samples of the two varieties of wheat studied.

