## Supplementary figures and images for "Infestation and growth of *Tribolium confusum* (Coleoptera: Tenebrionidae) on two varieties of local durum wheat: nutritional value and varietal susceptibility"

### figure jpg

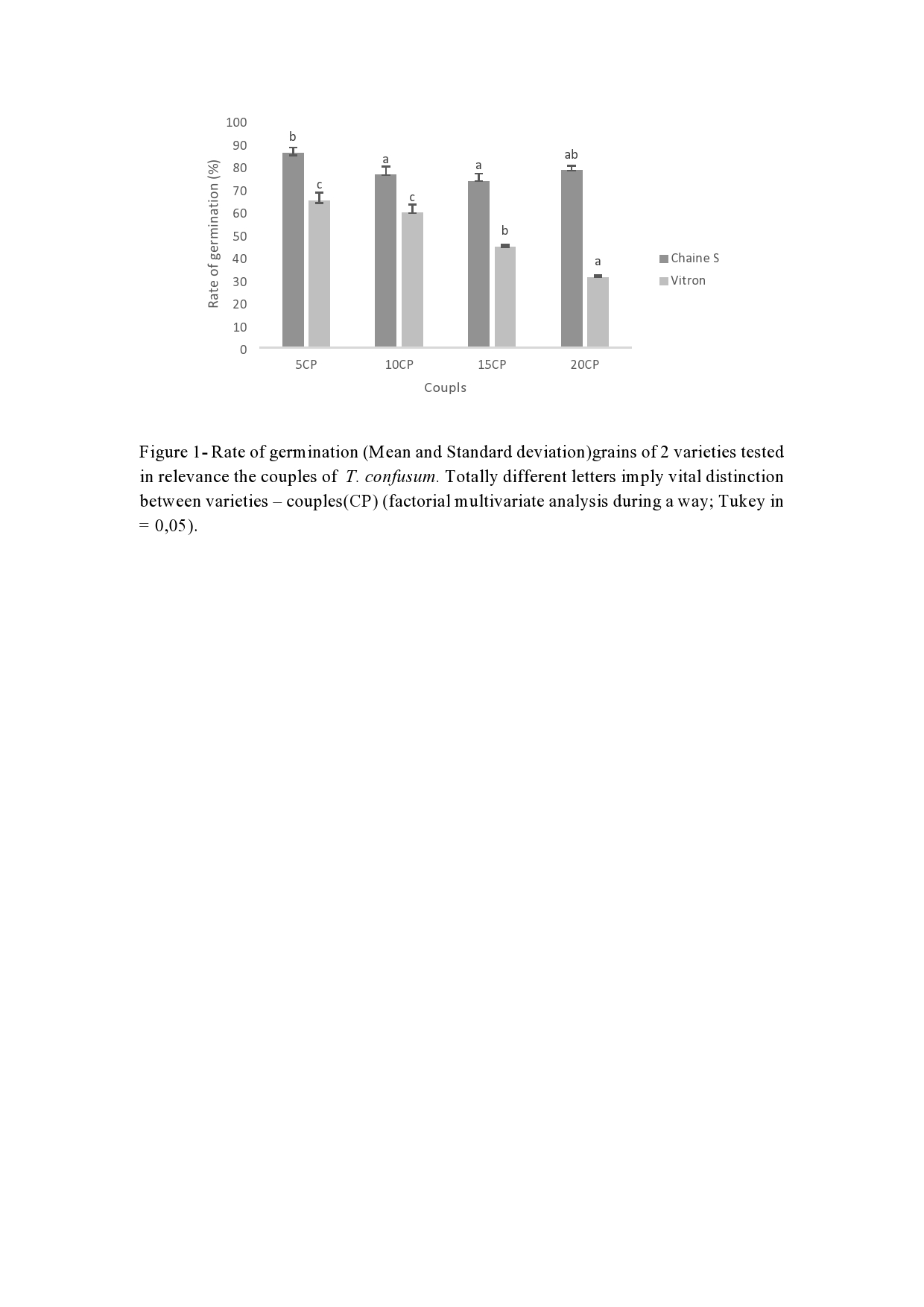
